# Evidences that host genetic background more than the environment shapes the microbiota of the snail *Bulinus truncatus*, an intermediate host of *Schistosoma* species

**DOI:** 10.1101/2024.11.01.619350

**Authors:** Mathilde J. Jaquet, Philippe Douchet, Eve Toulza, Thierry Lefevre, Bruno Senghor, Jérôme Boissier, Olivier Lepais, Emilie Chancerel, Benjamin Gourbal, Olivier Rey

## Abstract

Microbiota are increasingly recognized as key players in regulating host biological functions, influencing both the ecology and evolution of organisms. However, the factors shaping microbiota diversity and structure in natural environments remain underexplored, especially the relative importance of host genetics versus environmental factors. In this study, we address this gap using the freshwater snail *Bulinus truncatus*, an intermediate host for some human and animal *Schistosoma* parasites, as a model species. We developed 31 new microsatellite markers to assess the population structure of *B. truncatus* across 9 sites in Senegal. We then applied metabarcoding to characterize the diversity and structure of individual snail bacterial microbiota and environmental communities associated with each sampling site using environmental DNA. We also used molecular diagnostics to determine trematode infection status of *B. truncatus* individuals. By integrating these data through multiple regression on distance matrices (MRM) analyses, we quantified the influence of *B. truncatus* population genetics, spatial distribution, environmental bacterial communities, and infection status on the snail’s microbiota structure. Our results show that the genetic structure of *B. truncatus* populations, and to a lower extent geographic distribution, are the main factors explaining the snail’s microbiota compositions. Neither the environmental bacterial communities nor trematode infection status significantly contributed to microbiota structure. A portion of the variance in microbiota composition remains unexplained, suggesting that additional ecological or intrinsic factors might be involved. These findings provide new insights into the drivers of microbiota structure in natural populations and highlight the complexity of host-microbiota-environment interactions.

Key-words: *Bulinus truncatus*, Microbiota, Spatial structure, Population genetics, Trematodes, Multiple regressions on distance matrices

## 1. Introduction

The ‘microbiota’, consisting in all microorganisms associated with a host, is now broadly acknowledged as a crucial factor in various aspects of organismal biology, encompassing developmental, physiological, reproductive, behavioral and immune phenotypes (Liberti and Engel 2020; Daisley *et al*. 2020; Yuan *et al*. 2021; Davidson *et al*. 2024). The microbiota therefore partly influences the ecology and the evolution of their hosts and that of biotic interactions between host species (Sharon *et al*. 2010; Henry *et al*. 2021; Lange *et al*. 2023). Despite recent efforts to understand the factors shaping host microbiota, identifying and understanding the ecological and evolutionary determinants of microbiota composition and structure in natural host populations remains a major scientific challenge.

The composition and structure of natural hosts’ microbiota is particularly complex and depend on multiple environmental and host genetic factors (Benson *et al*. 2010). On one hand, the microbiota is partly composed of some either obligatory or facultative symbionts that are transmitted, generally via maternal inheritance, from one host generation to another (Funkhouser and Bordenstein 2013). Their evolutionary history hence follows that of their hosts with which they have co-evolved at both macroevolutionary and microevolutionary timescales leading to phylosymbiosis (Kohl 2020; Dapa *et al*. 2023). These microorganisms constitute the ‘core microbiota’ which consists of species consistently associated with the host regardless of environmental conditions (Risely 2020). Accordingly, the composition and structure of hosts’ microbiota is expected to be partly shaped by a combination of stochastic (e.g. dispersal, drift) and deterministic (e.g. selection) processes associated with hosts ecology and evolution (Benson *et al*. 2010; Furman *et al*. 2020; Hayashi *et al*. 2024).

On the other hand, most microorganisms that constitute hosts’ microbiota are facultative and are acquired from the environment throughout the host’s lifelong development. This is well illustrated by some species of Lepidopteran that lack a resident microbiota and generally acquire bacterial communities from their host plants or from the environmental microbial communities (Montagna *et al*. 2016; Phalnikar *et al*. 2018; Minard *et al*. 2019; Liu *et al*. 2020). Moreover, several environmental biotic factors including the communities of free-living microorganisms present in the environment (Díaz-Sánchez *et al*. 2018), and abiotic factors (e.g. temperature, pH) can influence the composition of the hosts’ microbiota which can in turn modify hosts biology (Bernardo-Cravo *et al*. 2020). In other word, the ecology of hosts (e.g. behavior, diet) also influences their microbiota (Archie and Tung 2015; Kennedy *et al*. 2020). Thus, it is expected that organisms from a common genetic pool and established in different environments harbor different communities of microorganisms (Berg *et al*. 2016). Accordingly, part of hosts’ microbiota is also expected to be partly shaped by ecological stochastic and deterministic (e.g. niche-based) processes (Hayashi *et al*. 2024). This more flexible microbiota compartment which includes taxa that are stable or predictable over time within individuals or across populations and is called the ‘temporal/dynamic core microbiota’ (Risely 2020).This more flexible microbiota compartment which includes taxa that are stable or predictable over time within individuals or across populations and is called the ‘temporal/dynamic core microbiota’ (Risely 2020). Our comprehension of factors shaping hosts’ microbiota hence requires accounting for the ecological context in which hosts are established and the genetic background of host populations.

Surprisingly however, few studies have yet specifically investigated the relative contribution of organisms’ genetic diversity and that of their environment in shaping natural hosts’ microbiota and lead to distinct conclusions. For instance, Suzuki and collaborators found that the structure of gut microbiota of wild house mouse (*Mus musculus domesticus*) populations is well predicted by genetic distances between host populations computed from an extensive exome genomic dataset and not by different environmental conditions, including temperature and diet (Suzuki *et al*. 2019). Conversely, the structure of the gut microbiota associated with California voles (*Microtus californicus*) populations across a contact zone between two recently diverged lineages of this species was best explained by the spatial distribution of hosts and not by lineage divergence, hence suggesting a strong influence of the environment on the structure of voles’ gut microbiota(Lin *et al*. 2020). In the same vein, Rothschild *et al*. (2018) found that the genetic background among 1046 healthy human individuals have minor role in determining individuals’ gut microbiota while environmental factors such as housing, diet and anthropometric measurements can explain some of the inter-individual variability in microbiota (Rothschild *et al*. 2018). Finally, in the *Nematostella vectensis* sea anemone model, Fraune et al. (2016) have shown that both the environment and genetics play a role in shaping the microbiota, and that these elements are complexly interconnected (Fraune *et al*. 2016). In the light of these studies, it appears that the importance of the genetic and environmental determinants of microbiota are potentially species and/or context dependent.

Here, we took advantage of high-throughput sequencing technologies to identify and assess the relative contribution of the ecological and host genetic factors structuring the microbiota associated with natural populations of the freshwater snail *Bulinus truncatus* as a model. *Bulinus truncatus* constitutes the main intermediate host for several trematode species including *Schistosoma haematobium*, the parasite responsible for urogenital bilharziasis in human populations in Africa (Toledo and Fried 2014). Ecological studies aimed at characterizing the communities of microorganisms associated with natural snail populations have recently flourished, especially in freshwater snails that are associated with disease transmission (Li *et al*. 2023). Based on cultivation-independent molecular methods, Van horn *et al*. (2012) have shown highly diverse intestinal microbial communities among three freshwater planorbid snails collected from the field and two of which serve has intermediate hosts of several parasites including digenetic trematodes of the genus *Schistosoma* the etiologic agent of Schistosomiasis (Van Horn *et al*. 2012). Similarly, Huot et *al.* (2020) demonstrated that seven species and strains of Planorbidae exhibited highly specific core bacterial composition that are closely related to the host strain. This bacterial composition show high congruence with the host phylogeny, hence revealing a phylosymbiotic pattern (Huot *et al*. 2020). More recently, (McCann *et al*. 2024), showed that the microbiota of *Galba truncatula*, the main intermediate host for the zoonotic trematode *Fasciola hepatica*, vary between natural populations established at two geographically and ecologically distinct sites, hence highlighting the existence of natural diversity in the composition of bacterial communities associated with this snail species. However, whether such microbiota diversity in natural populations is driven by snails’ genetic background or by environmental factors remains to be fully elucidated. If the freshwater snails’ microbiota is important in the circulation of associated pathogens, as has been suggested, it is essential to study the natural diversity, the structure and the factors structuring these communities in the field.

Here, we characterized the diversity and structure of the whole-body *B. truncatus* microbiota over 9 freshwater aquatic sites in Northern Senegal. The presence of trematodes developing within each snail was molecularly diagnosed and the genetic background of each snail host was characterized based on 31 newly developed microsatellite markers using a genotyping-by-sequencing approach. Aquatic bacterial communities associated with each of the 9 studied freshwater sites were also characterized and used as a proxy of the ecological context in which *B. truncatus* populations are established. We used these complementary datasets to identify the main factors structuring the microbiota associated with *B. truncatus* in natural populations.

## 2. Materials and Methods

### 2.1. Field collection of samples

Field sampling was conducted in February 2022 during the dry season in the Senegal River basin region (**Fig. 1**). We focused on nine natural sites of *S. haematobium* transmission, where urogenital schistosomiasis’ prevalence among school-aged children was previously reported (Senghor *et al*. 2022). These sites differ in terms of ecological contexts and in terms of *B. truncatus* abundance (Douchet *et al*. 2024). Six transmission sites are located near the villages of Ndiawara, Ouali Diala, Dioundou, Fonde Ass and Khodit, all of which being located along river “le Doue”, a tributary of the Senegal River in the middle valley. One transmission site, near the village of Guia, is also located in the middle valley and consists of an irrigation canal that drains water from the river “le Doue”. Two transmission sites, near the villages of Mbane and Saneinte are located along the east shore of Guiers lake. One transmission site near the village of Lampsar is located in the lower valley and consists of an inlet of the Senegal River delta (**suppl. Table 1**).

**Figure 1:**
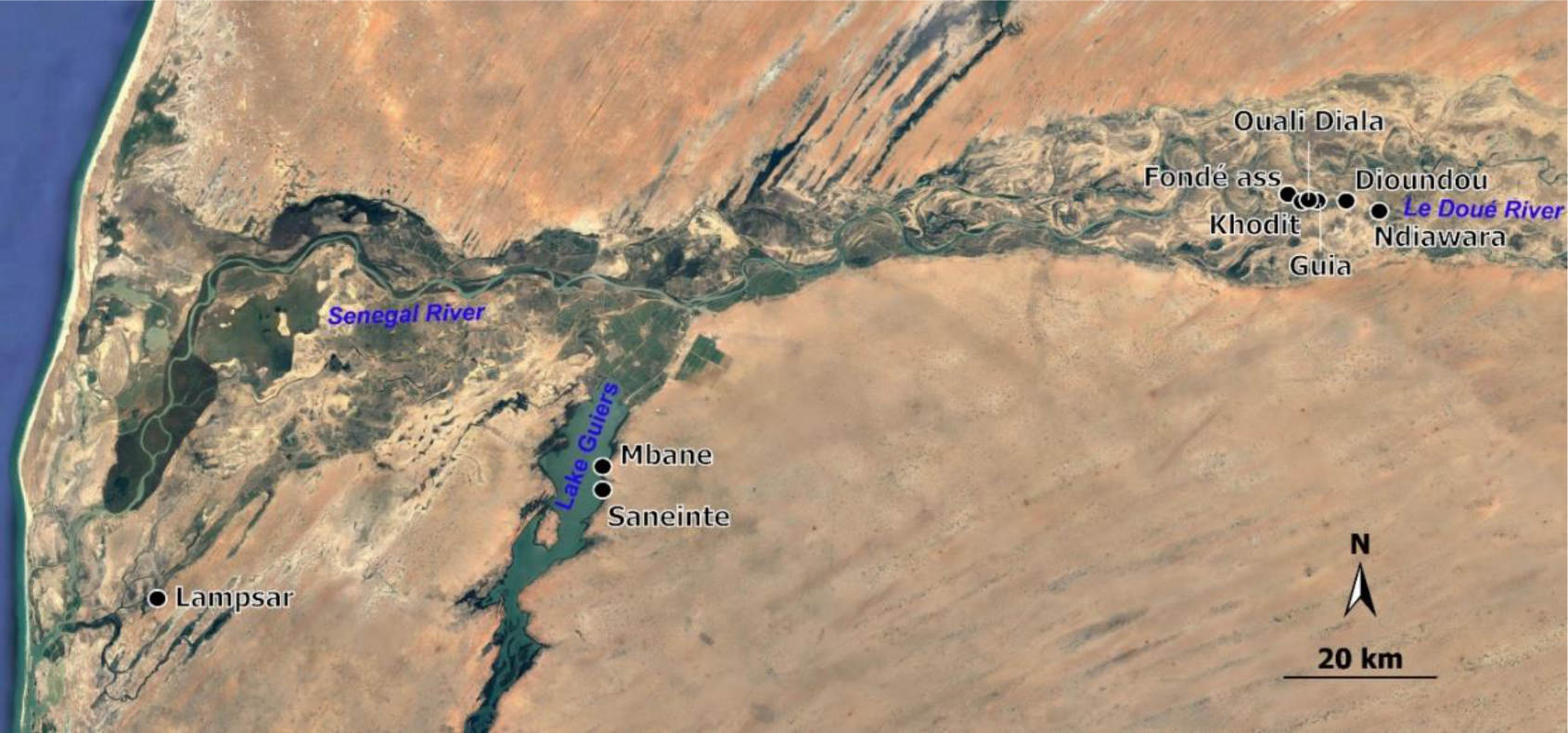
Locations of the studied aquatic transmission sites (black dots) nearby the closest established villages, on a satellite map of northern Senegal.

#### 2.1.1 Field collection of environmental samples

At each site, we first filtered water along the water column from the surface to the water-sediment interface. We used filtration capsules of 0.45 µM mesh size (Waterra USA Inc.) connected to an electric water pump as previously described in Douchet and collaborators (Douchet *et al*. 2022). Filtration was performed until the filtration capsule was clogged and the final filtration volume was retrieved. Once the filtration completed, the filtration capsule was depleted from its water content, filled with 50 mL of Longmire solution, vigorously shaken, and preserved at ambient temperature until used for environmental DNA (eDNA) extractions. At each site but Khodit, a technical field negative control was obtained by filtering 1.5L of commercial spring water, following the same protocol for preservation.

#### 2.1.2 Field collection and preservation of snails’ samples

Following eDNA sampling, we manually collected all encountered aquatic snails by scooping the aquatic vegetation using a colander for about 30 minutes to 1 hour, along a 10 to 30-meters long transect along the shore of the targeted waterbody (either a river or a lake). After taxonomic identification (following the keys of (Mandahl-Barth 1962)), up to 17 non-emitting *B. truncatus* snail were extracted from their shell, after a quick heating step at 70°C for 1 minute, with decontaminated forceps and individualized into a sterile 1.5ml tube containing 70% Ethanol. Forceps were decontaminated between each sample using a DNA AWAY solution. The extraction of snail bodies from their shells was achieved to avoid DNA contamination with microorganisms, including bacteria, established on snail shells. Overall, 124 (mean = 14; min = 5; max = 17; **suppl. Table 1**) adult *B. truncatus* snails that displayed a shell size of approximately 6 mm long, were prepared as above.

### 2.2. DNA Extraction from eDNA samples and *B. truncatus* snails

All pre-PCR steps including DNA extraction and the preparation of PCR mixes were conducted under a sterile hood decontaminated before and after each use with 10% bleach, 70% ethanol, a DNA AWAY solution followed by a UV light exposure for 20 minutes.

Total eDNA from water filtrations were extracted following (Douchet et al. 2022). Briefly, the Longmire buffer contained within each filtration capsule was equally split into three 50 mL tubes. For the field negative controls (i.e. Spring water filtrates), each capsule content was recovered in one single 50 ml tube. The tubes were centrifuged for 20 min at 16,000 x g and the supernatant was discarded. We then collected up to 1g of sediment from the pellet from each tube or 500 µl of Longmire remaining at the base of the tube when not enough material was observed. Thus, total eDNA from each capsule were extracted in triplicate. No pellets were observed for filtration negative controls (consisting in spring water) after centrifugation. For these samples, 500 µL of Longmire were retained and resuspended after removing the supernatant and were processed as the other samples. This step led to the processing of 35 samples (i.e. 3 extraction replicates for each of the nine eDNA samples and one field negative control per site except for Khodit). The total eDNA of these 35 samples was extracted using the DNeasy PowerSoil Pro kit (Qiagen) according to manufacturer’s protocol performing the physical lysis with a MagNA Lyser at a speed of 7000 × g for 30 s.

Total genomic DNA from each of the 124 snails removed from their shell was extracted using DNeasy 96 Blood and Tissue Kit (Qiagen), according to the manufacturer’s protocol. Two negative control extractions consisting of empty 1.5 ml tubes same as those used for snail preservation were added in the process.

### 2.3. Infection status and genotyping of *B. truncatus* snails

#### 2.3.1. Infection status of *B. truncatus* snails

The presence of developing trematodes within each of the 124 snails was diagnosed using the Trem_16S_F1 (GACGGAAAGACCCCRAGA) and Trem-16S_R2 (CRCCGGTYTTAACTCARYTCAT) 16S trematode metabarcode (Douchet *et al*. 2022). DNA extracts were diluted to 1/100^th^ to minimize PCR inhibitors and hence limit PCR false negatives. PCR reactions were prepared and run under standard conditions, with detailed methodologies provided in **Appendix 1**. Positive samples were reamplified and sent to the Bio-Environment platform (University of Perpignan Via Domitia, France) to identify the trematodes potentially developing within snail hosts and accounting for possible coinfections. Libraries were performed using Illumina Nextera index kit and Q5 high fidelity DNA polymerase (New England Biolabs). Indexed PCR products were then normalized with SequalPrep plates (ThermoFisher) and paired-end sequenced on a MiSeq instrument using v2 chemistry (2 x 250 bp).

#### 2.3.2. Microsatellite development

The *B. truncatus* genome (GenBank accession GCA_021962125.1, (Young *et al*. 2022)) was used for microsatellite discovery, leading to the selection of 96 primer pairs optimized for multiplex PCR (details in **Appendix 2**).

#### 2.3.3. Sequence-based microsatellite genotyping

Multiplex PCR amplification and sequencing libraries were constructed, with detailed methods provided in **Appendix 3**.

#### 2.3.4. Genetic analyses

Descriptive statistics of genetic diversity including the mean number of alleles, the mean allelic richness (computed based on the minimal number of genotypes obtained over sampling sites, N = 5), the mean expected (He) and observed (Ho) heterozygosity, and the mean F*is* were computed at each sampling sites using the SPAGeDi software (v. 1.5d; (Hardy and Vekemans 2002)) accounting for the allotetraploidy nature of *B. truncatus* (Njiokou *et al*. 1993).

We next computed pairwise Nei’s genetic standard distance Ds (Nei 1978) and F_st_ values between each sampling sites using the SPAGeDi software (v. 1.5d; (Hardy and Vekemans 2002)). From the obtained Ds values, we ran a Mantel test to test for possible isolation by distance pattern among sampling sites using matrices of pairwise Ds values and pairwise linear geographical distances obtained between each sampling site. The statistical significance of the Mantel coefficient r was tested based on 10000 permutations as implemented in the ‘*adegenet’* R package (Jombart 2008). A similar Mantel test analysis was run using pairwise (F*_st_* / (1 - F*_st_*) values obtained between sampling sites.

### 2.4. Sequencing of bacterial communities and metabarcoding analyses

A total of 198 bacterial *16S* metabarcoding libraries were prepared following the Illumina two-step PCR protocol; including the DNA extracts of the 124 snails, 70 eDNA replicates consisting in 54 eDNA samples (each extraction triplicate was amplified in duplicate) and 16 technical field negative controls (each control was amplified in duplicate), and 4 negative PCR controls (milliρ water) (**suppl. Table 3**). We targeted the variable V3-V4 loops region of the *16S* sDNA gene using the 341F (5’-CCTACGGGNGGCWGCAG-3’) and 805R (5’-GACTACHVGGGTATCTAATCC-3’) primers (Klindworth *et al*. 2013) combined with universal Illumina adapters.

The first PCRs were performed using the Q5® High-Fidelity 2X Master Mix (New England BioLabs), in a 25µL final volume containing 2 µL of template DNA and using a PCR program consisting in an initial denaturation step of 30s at 98°C followed by 32 cycles containing a denaturation step of 6 sec at 98°C, an annealing step of 30 sec at 55°C, and an elongation step of 8 sec at 72°C and ending with a final elongation step of 60 sec at 65°C. We used 5 µL of the PCR products to check the PCR products quality and integrity through electrophoresis on agarose gels. The 20µL left were sent to the Bio-Environment platform (University of Perpignan Via Domitia, France). Libraries were performed using Illumina Nextera index kit and Q5 high fidelity DNA polymerase (New England Biolabs). Indexed PCR products were then normalized with SequalPrep plates (ThermoFisher) and paired-end sequenced on a MiSeq instrument using v2 chemistry (2 x 250 bp).

Raw data were processed using R (version 2023.03.0+386) using a pipeline based on the ‘*dada2*’ package (v. 1.28.0) (Callahan *et al*. 2016) on the IFB (French Institute of Bioinformatics) cloud. Briefly, this pipeline consists in removing primers from the obtained sequenced reads (max.mismatch = 1), trimming and quality filtering reads (minLen = 150, maxN = 0, maxEE = c(3, 3), truncQ = 2), denoising, dereplicating, merging paired-end reads (maxMismatch = 0), building the Amplicon Sequence Variant (ASV) table and removing potential chimeric sequences produced during the process (method = ‘consensus’). The taxonomic assignment (multithread = TRUE, minBoot = 60) of the resulting filtered and cleaned ASVs was performed using the Ribosomal Database Project (RDP) classifier (Wang *et al*. 2007) and based on the Silva_train_set and silva_species_assignment 138.1 datasets (Quast *et al*. 2012). Eukaryotic, mitochondrial, and chloroplast sequences were removed, as well as ASVs not affiliated to ‘Bacteria’ at the Kingdom taxonomic levels. The obtained ASV sequence table was finally merged with the taxonomy database and the sample metadata matrix for subsequent analyses using the ‘*phyloseq’* R package (v. 1.22.3) (McMurdie and Holmes 2013).

The final ASV dataset was next cleaned from possible contamination during field collection and biomolecular processing (DNA extraction, PCR). To this aim, ASVs obtained from negative PCR controls were removed from all samples through the ‘prune_taxa’ function of the ‘*phyloseq*’ package. Then, the ‘merge_phyloseq’ function of the ‘*phyloseq*’ package was used to generate the whole object containing all the samples. Moreover, ASV present at low abundance (< 0.005%; (Bokulich *et al*. 2013)) were filtered from the whole dataset to account for possible sequencing errors. Briefly, the whole phyloseq object was transformed in relative abundance with the ‘transform_sample_count’ function of the ‘*phyloseq*’ R package. Then, the low abundant taxa were removed through the ‘prune_taxa’ function of the ‘*phyloseq*’ R package.

To assess whether sequencing depth was satisfactory across samples, rarefaction curves were generated using ‘rarecurve’ function from the ‘*vegan*’ R package (v. 2.6.4) (Oksanen *et al*. 2022). The abundance of each ASV was normalized based on the lowest sample in terms of ASV counts (i.e. 7000, see **suppl. Table 3**) by random sub-sampling using the ‘rarefy_even_depth’ function of the ‘*phyloseq*’ package (rngseed = 1000) (McMurdie and Holmes 2013). To quantify read counts per sample during preprocessing steps, refer to **suppl. Table 3**.

### 2.5. Statistical analyses

#### 2.5.1. Diversity in environmental and snail-associated bacterial communities and characterization of *B. truncatus* core microbiota

The diversity of bacterial communities associated with environmental matrices and *B. truncatus* snails was assessed at the sampling site level based on the absolute number of ASVs, the Shannon and Simpson’s diversity indices as implemented in the ‘*microbiome*’ R package (v. 1.22.0) (Lahti and Shetty 2012). Kruskal-Wallis nonparametric tests were computed to compare each of these alpha-diversity indices between sample type (i.e. snails *versus* eDNA samples) and between sites using the ‘*stats*’ R package (v. 4.3.0) (R Core Team 2023). Those tests were followed by pairwise multiple comparisons using the ‘FSA::dunnTest’ of the ‘FSA’ R package (v. 0.9.5) with the Benjamini-Hochberg method for adjusting p-values (Dunn 1964).

To characterize the core microbiota of *B. truncatus in natura,* which consists in the most stable part of the bacterial communities shared among all individuals (Risely 2020), we followed established guidelines (Neu *et al*. 2021). The core microbiota was assessed at the family level using the ‘core_members’ function of the ‘*microbiome*’ R package (Lahti and Shetty 2012) and setting the detection parameter to 0 and the prevalence parameter to 90%.

#### 2.5.2. Identification of factors structuring environmental and snail bacterial communities

To investigate for possible structuration of bacterial communities in environmental samples and *B. truncatus* snails we conducted a Principal Coordinate Analysis (PCoA) using the ‘pco’ function of the ‘*ecodist*’ package (v. 2.1.3) (Goslee and Urban 2007). Pairwise Jaccard and Bray-Curtis indices were conducted using the ‘distance’ function of the ‘*phyloseq*’ R package. Based on these beta diversity indices, we ran permutational multivariate analyses of variance (PERMANOVA) to test for: 1-Differences between aquatic bacterial communities (i.e. eDNA samples) and snails associated communities, accounting for the sampling site as an additional explanatory variable and for a possible interaction between sampling site and the nature of the sample (environment versus snail). 2-An effect of trematode infection status (all trematode species combined) on the difference in *B. truncatus* microbiota. Here we accounted for the sampling site as an additional explanatory variable and an interaction term between the sampling site and the infection status. PERMANOVA were conducted using the ‘adonis2’ function implemented in the ‘*vegan*’ R package (Oksanen *et al*. 2022), and setting the number of permutations to 10 000.

To assess the relative effect of the environmental bacterial communities and the genetic background of snails on whole-body snail microbiota, we tested for correlation between four pairwise matrices including, the geographical distance matrix between each sampling site, the Jaccard indices computed between environmental bacterial communities at each sampling site, the Jaccard indices computed between bacterial communities associated with *B. truncatus* individuals and the Nei’s genetic distance between snail hosts. To ensure comparable dimensions for the four matrices, 14 samples from the original matrices were discarded using the ‘subset_samples’ function from the ‘*phyloseq*’ R package. This step led to two matrices of 109*109 dimensions. Based on those matrices, we pseudoreplicated the geographic distance and the Jaccard indices computed between environmental bacterial communities at each sampling site matrices to transform them in the same dimensions.

Based on these four matrices, we conducted a multiple regression on distance matrices (MRM) fixing the Jaccard distance matrix obtained from the bacterial communities between *B. truncatus* individuals as the dependent variable and using three matrices as explanatory variables: the Jaccard distance between bacterial communities at each site (pseudoreplicated at the individual level), the Nei’s genetic distance computed between individuals and the euclidean geographic distance between sites (also pseudoreplicated at the individual level). This MRM analysis was ran using the ‘MRM’ function of the ‘*ecodist*’ R package and the significance of this multiple regression test was assessed based on 9999 permutations.

Results were visually assessed using a custom function to plot matrices against one another, with Mantel test statistics computed using the ‘mantel’ function of the ‘*vegan*’ R package.

## 3. Results

### 3.1. Abundance of *B. truncatus*, trematode prevalence among sites

Overall, from the 124 *B. truncatus* extracted from their shell, the total trematode infection prevalence was 15.3% ranging from 0% (Dioundou, Mbane, Saneinte) to 29.4% (Fonde Ass) (**suppl. Table 1)**. Nine species of trematodes were identified. Aside from *Schistosoma haematobium* and *Schistosoma bovis,* we also found *Haematolochus* sp., two species of Paramphistomoidea, *Orientocreadium batrachoides*, *Petasiger* sp. and two species of Diplostomoidea.

### 3.2. Genetic diversity and structure of *B. truncatus* among sampling sites

Based on the 31 newly developed microsatellite markers, we found little genetic diversity with little variation in genetic diversity among sites. Allelic richness (Ar) ranged from 2.13 (Dioundou) to 2.75 (Mbane) and observed heterozygosity (Ho) ranged from 0.38 (Ndiawara and Saneinte) to 0.41 (Guia and Mbane) (**Table 1**). Inbreeding coefficients (F_is_) were different from zero for all but the two sites Mbane and Saneinte located on the Guiers lake. Pairwise Ds genetic distances computed between sites ranged from 0.0019 between Ndiawara and Dioundou both sites being located along the Doué river approximately 4.8 km one from each other; and 0.4354 between Ouali Diala (Doué River) and Lampsar distant by approximately 160.3 km. The observed genetic differentiation between sites followed an isolation by distance pattern (Mantel r = 0.76; Pval = 0.0009) (**suppl. Fig. 1**).

**Table 1.** Genetic diversity statistics for *B. truncatus* at each sampling site based on the 31 microsatellite markers developed in this study. Na: Number of alleles; Ar: Allelic richness; He: gene diversity (Nei 1978); Ho: Observed heterozygosity; Fis: Inbreeding coefficient; Pval of Fis after 10000 randomization of gene copies among individuals.

| Site | Sample size | Na | Ar | He | Ho | Fis | Pval (Fis<>0) |
| --- | --- | --- | --- | --- | --- | --- | --- |
| Ndiawara | 7 | 2.52 | 2.42 | 0.405 | 0.382 | 0.063 | 0.028 |
| Ouali Diala | 11 | 2.42 | 2.27 | 0.375 | 0.389 | -0.042 | 0.054 |
| Guia | 16 | 2.58 | 2.21 | 0.384 | 0.406 | -0.06 | 0.001 |
| Dioundou | 5 | 2.13 | 2.13 | 0.369 | 0.396 | -0.087 | 0.011 |
| Fonde Ass | 15 | 3.06 | 2.53 | 0.417 | 0.39 | 0.069 | 0.001 |
| Khodit | 15 | 2.97 | 2.63 | 0.453 | 0.396 | 0.133 | 0 |
| Mbane | 12 | 3.23 | 2.75 | 0.401 | 0.41 | -0.023 | 0.239 |
| Saneinte | 16 | 3.26 | 2.68 | 0.383 | 0.382 | 0.004 | 0.806 |
| Lampsar | 13 | 2.9 | 2.57 | 0.421 | 0.392 | 0.072 | 0 |
| Overall sites | 110 | 5.52 | 3.18 | 0.499 | 0.394 | 0.211 | 0 |

### 3.3. Diversity in environmental and snail-associated bacterial communities and characterization of *B. truncatus* snails’ core microbiota

In terms of alpha diversity, the bacterial communities from the environment displayed equal alpha diversity among sites except for Lampsar that displayed lower alpha diversity whatever the index used (i.e. Species Richness, Shannon, Simpson) (**suppl. Fig. 2**). Moreover, *B. truncatus* microbiota displayed lower diversity than aquatic bacterial communities, except for the Lampsar site, when considering the ASV richness (Kruskall-Wallis chi squared = 95.394; Pval < 0.0001), the Shannon diversity index (Kruskall-Wallis chi squared = 92.187; Pval < 0.0001) and the Simpson index (Kruskall-Wallis chi squared = 89.132; Pval < 0.0001) (**suppl. Fig. 2 and suppl. Table 4)**. We found significant differences in the mean alpha diversity indices in the microbiota of *B. truncatus* from the different sample sites (all p < 0.05, **suppl. Table 5** and **suppl. Fig. 3A**). Pairwise post-hoc tests revealed that only the alpha diversity of the sampling site Mbane was lower than that of Saneinte ((Z = 6.78, Pval < 0.01), Guia (Z = 3.16, Pval = 0.014) and Lampsar (Z = 8.17, Pval < 0.01). Moreover, we found no significant differences in alpha diversity between infected and uninfected snails (all p > 0.05, **suppl. Table 6 and suppl. Fig. 3B**).

Proteobacteria were predominant in the microbiota of *B. truncatus* across all sampling sites, followed by Bacteroidota and Firmicutes (**Fig. 2A**). Among the phyla identified in the environmental samples, only two were common to both the snail microbiota and the environmental samples: Cyanobacteria and Planctomycetota (**suppl. Fig 4**). Cyanobacteria was the second most abundant phylum in the environmental samples, while Planctomycetota ranked seventh among the 28 phyla found. Interestingly, Planctomycetota were more abundant in the snail samples than in the environmental samples, whereas the opposite was true for Cyanobacteria.

**Figure 2:**
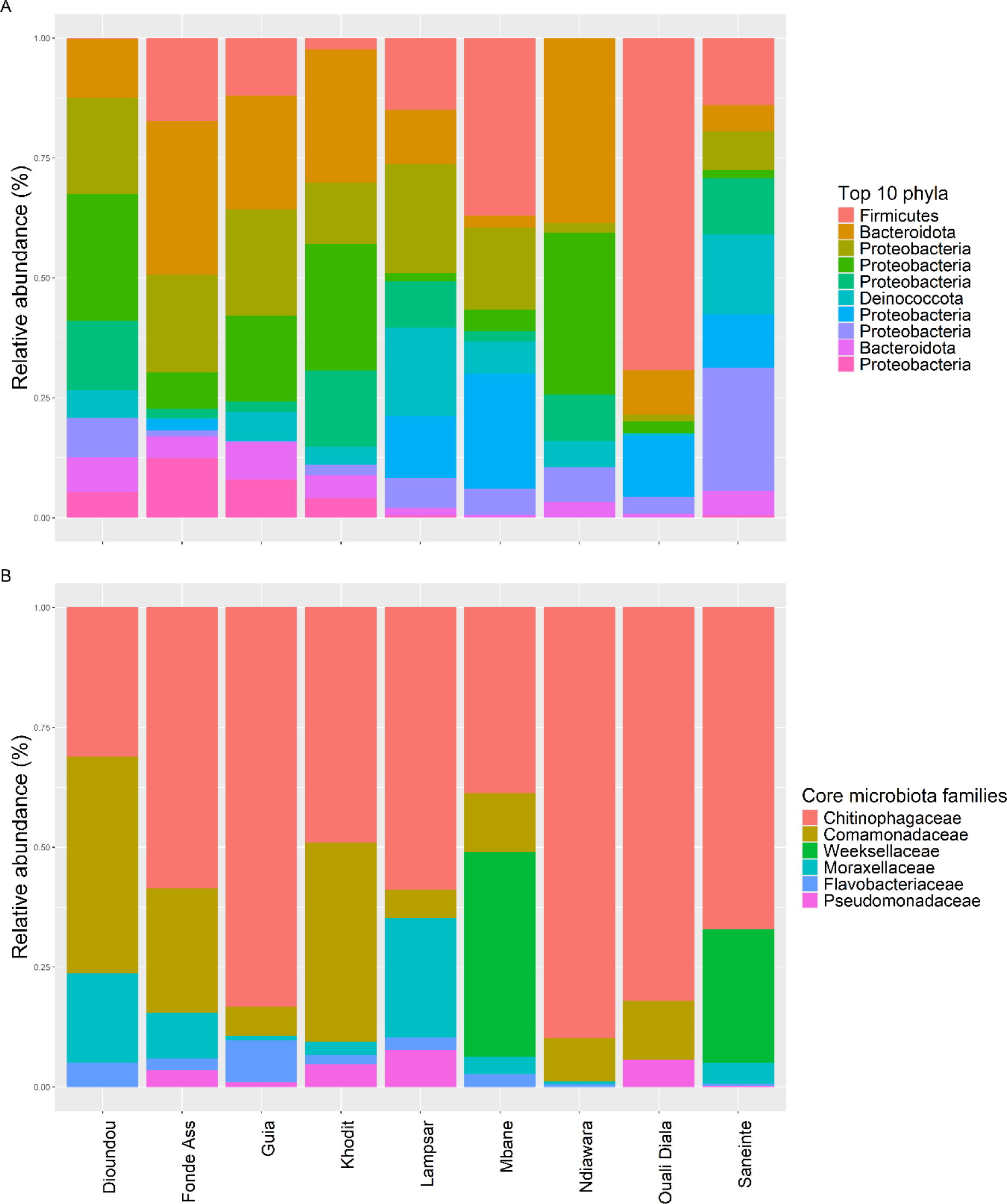
(A) Relative abundance of the ten most abundant phyla in the microbiota of *B. truncatus* from the nine different sites. (B) Bacterial composition of the core microbiota of *B. truncatus* snails at the family level across the 9 studied sites.

The core microbiota of *B. truncatus* was characterized at the family level by considering bacterial families present in at least 90% of the overall snails. The core microbiota consisted of 6 families out of the 169 families identified in the snails. These families, which included Chitinophagaceae (69.3% of core microbiota reads), Weeksellaceae (12.9%), Comamonadaceae (8.9%), Moraxellaceae (3.4%), Flavobacteriaceae (2.9%), and Pseudomonadaceae (2.6%), were common to at least 90% of individuals across all sampling sites, although their relative abundances varied (**Fig. 2B**).

### 3.4. Identification of factors structuring environmental and snail bacterial communities

#### 3.4.1. The microbiota of *B. truncatus* snails differ from environmental bacterial communities

Based on the microbial communities characterized from the 54 environmental samples and the 124 *B. truncatus* individuals, we found that the total microbiota associated with whole-body *B. truncatus* differs from bacterial communities present in the environment. The first two axes of the Principal Coordinates Analysis (PCoA) performed on the Jaccard distance matrix obtained among all samples (i.e. *B. truncatus* snails and environmental samples) and all sampling sites explained 21.83% of the total variance (**Fig. 3**). *Bulinus truncatus* and environmental samples cluster into two distinct groups along these two axes. Such difference in the composition in bacterial communities between the environmental samples and *B. truncatus* is confirmed by the PERMANOVA analysis which indicates significant effect of the type of sample (i.e. *B. truncatus* snail and environmental sample) (F-value = 39.44; Pval < 0.0001) and of the sampling site (F-value = 6.42; Pval < 0.0001). Similar results were obtained from the PERMANOVA analysis based on the Bray-Curtis distances (nature of the sample: F-value = 76.60; Pval < 0.0001; site: F-value = 10.66; Pval < 0.0001; **suppl. Fig. 5**).

**Figure 3.**
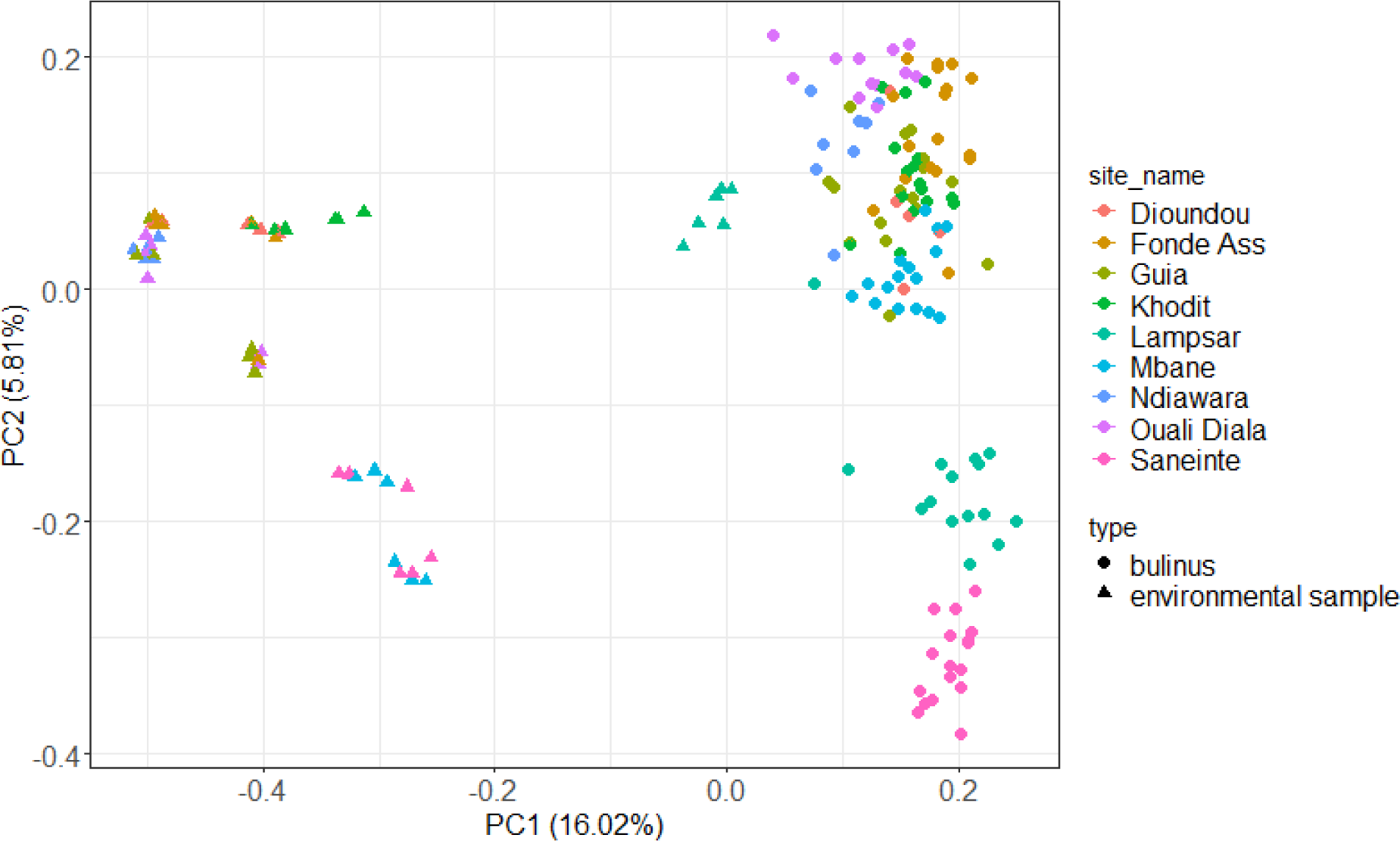
Principal coordinate analysis (PCoA) plot showing environmental samples (triangles) and *B. truncatus* snail associated (dots) bacterial communities distributed along the two first PCoA axes computed based on pairwise Jaccard distances obtained between each pair of samples.

#### 3.4.2. The microbiota of *B. truncatus* snails is geographically structured, partly driven by their genetic background and the geographic distance and to a lesser extent by the environment

The first two axes of the Principal Coordinates Analysis (PCoA) performed based on the Jaccard distance matrix obtained from *B. truncatus* microbiota explained 28.36% of the total variance (**Fig. 4**). Our PERMANOVA analysis supports an effect of the sampling site with the structure of *B. truncatus* microbiota (F-value = 7.84 Pval < 0.0001) (**Fig. 4**). The same results were observed using pairwise Bray-Curtis distances (F-value = 12.20; Pval < 0.0001, **suppl. Fig. 6**). Conversely, we found no effect of the infection status of *B. truncatus* on the beta diversity of snails’ microbiota, neither based on the Jaccard distances (**Fig. 4**) nor based on the Bray-Curtis distances (all p > 0.05, **suppl. Table 7** and **suppl. Fig. 6**) even when accounting for the effect of the sampling site.

**Figure 4:**
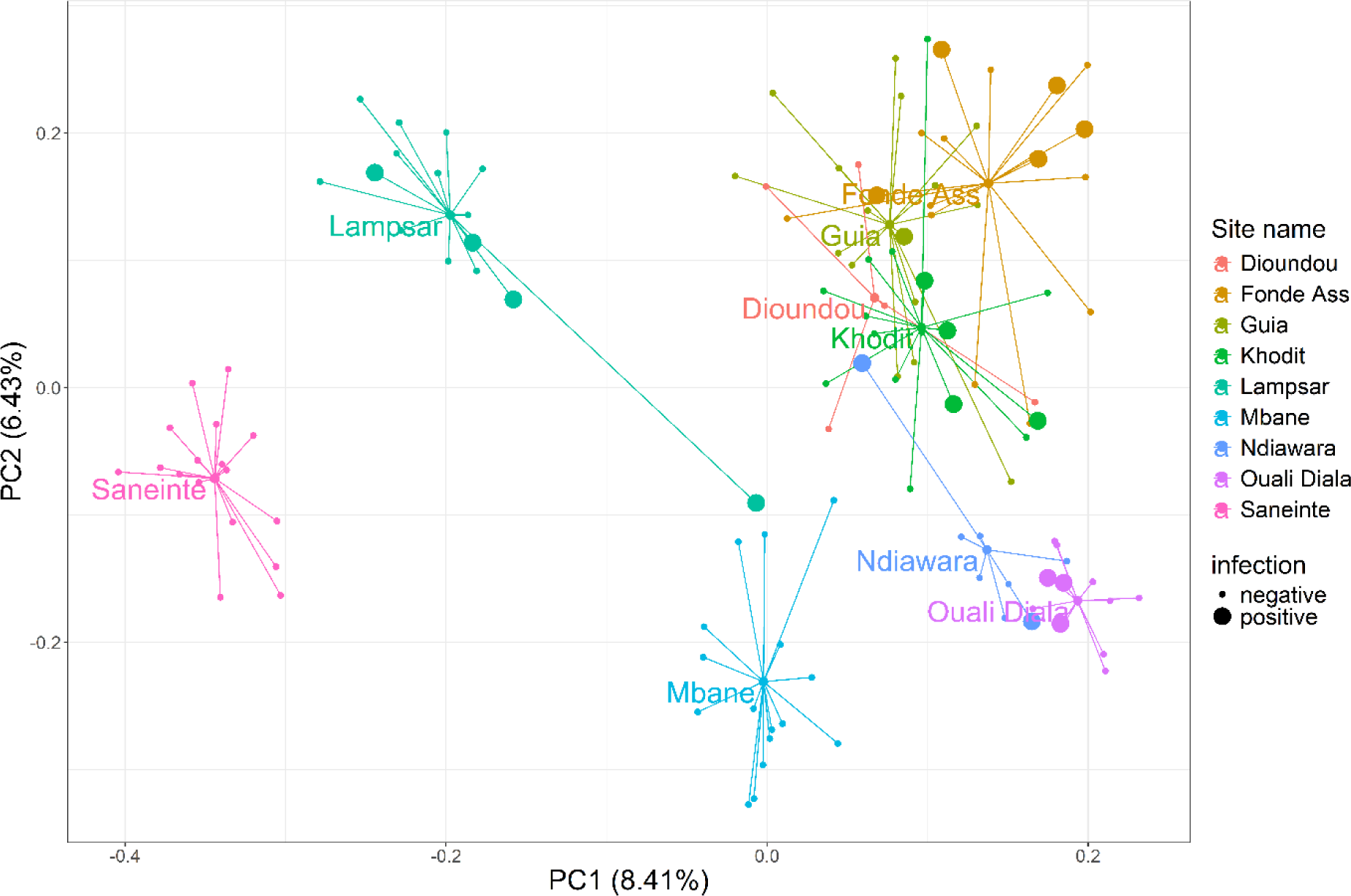
Principal coordinate analysis (PCoA) plot showing the bacterial communities of *B. truncatus* according to their original sampling sites (see color legend), and to their infection status by trematodes based on a molecular diagnostic (dot size) distributed along the two first PCoA axes based on pairwise Jaccard similarity indices computed between each sample. Each colored small dot represents one snail host from a given sampling site. Large dots are snail hosts positive to trematode molecular diagnostic. Medium dots are the centroids of the bacterial communities of snails originating from each sampling site.

Based on the multiple regression on distance matrices (MRM) and on our series of correlation tests (**Fig. 5**), the variables that explain the mean Jaccard indices calculated between *B. truncatus* microbiota at the individual level are, in order of importance, the Nei’s genetic distance between individuals (r = 0.24; Pval < 0.001), the geographical distance between sites (r = 0.22; Pval = 0.001) and the mean Jaccard indices calculated between the aquatic bacterial communities at the site level (r = −0.05; Pval = 0.39) both pseudopseudo-replicated at the individual level.

**Figure 5:**
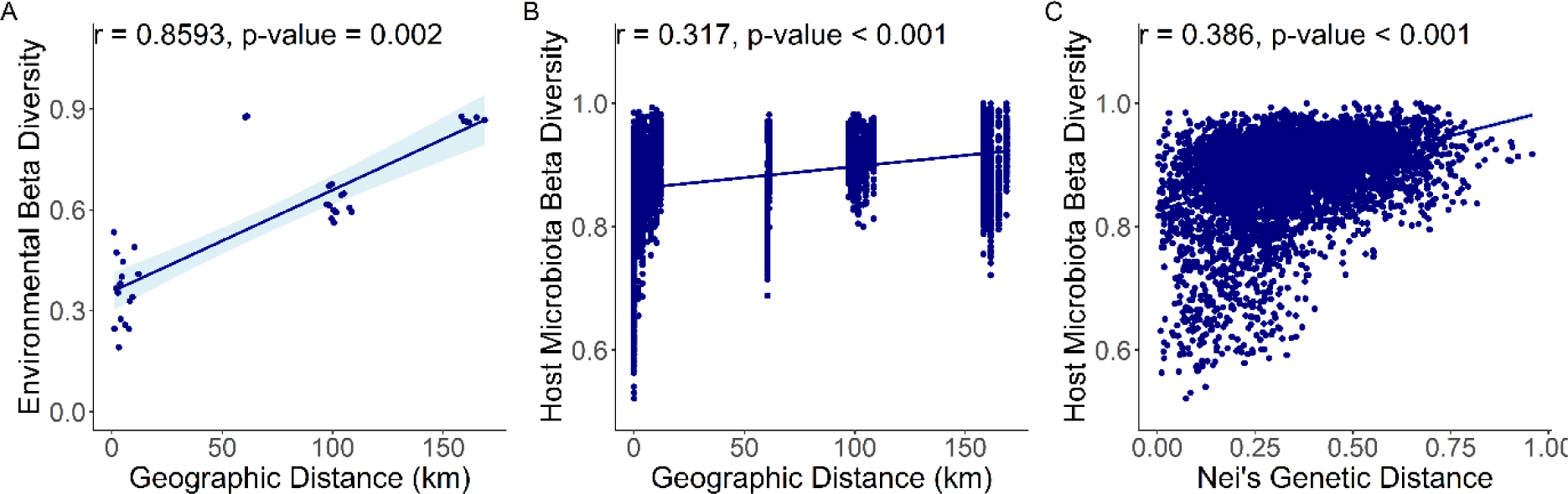
Distance-decay scatterplots showing bacterial communities distance (determined by the pairwise Jaccard similarity index) as a function of geographic distance across sampling sites for environmental samples (A), and *B. truncatus* snails (B). Scatterplot of *B. truncatus* snails microbiota dissimilarity (determined by the Jaccard index) as a function of Nei’s genetic distance between *B. truncatus* individuals computed based on 31 microsatellite loci (C). Blue lines of best fit are included to highlight trends.

## 4. Discussion

### 1. Composition of the microbiota in natural populations of *B. truncatus*

#### 1.a. Bacterial communities display lower alpha-diversity in *B. truncatus* than in environmental samples

Our results provide new insights on the composition of the microbiota associated with *B. truncatus* natural populations. Overall, *B. truncatus* display microbiota less diversified than bacterial communities present in the surrounding freshwater environment. This was somewhat expected because bacterial communities collected from the environment encompass free-living bacteria, bacteria that are released by all organisms associated with freshwater habitats and bacteria associated with freshwater microorganisms that could have been trapped during eDNA sampling (e.g. algae, invertebrates) (Mikhailov *et al*. 2019; Gendron *et al*. 2019; Sadeghi *et al*. 2021). Moreover, host microbiota are generally considered as a subsample of environmental bacterial communities resulting from a filter induced by the intrinsic properties of the hosts (Reese and Dunn 2018; Mazel *et al*. 2018; Suzzi *et al*. 2023). In particular, the host constitutes an ecological niche, which suitability for bacteria is largely influenced by hosts immune system that acts as a selective pressure, promoting the growth of some beneficial bacteria while inhibiting potential pathogens and maintaining the microbiota homeostasis (Belkaid and Harrison 2017). Such lower alpha-diversity in host microbiota compared to environmental bacterial communities was also described in recent studies conducted on other freshwater snails including *Ampullaceana balthica* and *Galba truncatula* (Herlemann *et al*. 2024a; McCann *et al*. 2024).

We found only little inter-site differences in *B. truncatus* microbiota alpha diversity among the nine sampling sites. The microbiota of *B. truncatus* from only one site (i.e. Mbane) harbored lower alpha-diversity than two other sites (i.e. Saneinte and Lampsar). The low alpha diversity of *B. truncatus* microbiota observed in Mbane comparatively to those of Saneinte is intriguing since these two sites are geographically distant from only 3.2 km and established along the same shore of the Guiers Lake. In our study, no obvious technical (e.g. difference in sequencing coverage), or measured biological factors (e.g. host size, host genetic diversity), can satisfactorily explain this pattern either at an individual or site scale (data not shown). Ecological differences between these two sites, and in particular in terms of food resources available for local *B. truncatus* populations, could be one explanatory factor explaining such difference, although we did not collect this information in the field. Studies have suggested that, additionally to host physiological traits, dietary diversity (exogenous microbial diversity) also impact gut microbiota alpha diversity (Reese and Dunn 2018). Fine-scale geographical studies at the scale of the lake accounting for more detailed ecological factors cancan be useful to better understand such result.

#### 1.b. *Bulinus truncatus* microbiota composition in natural populations

We observed that Proteobacteria, Bacteroidota, and Firmicutes were the dominant phyla in the microbiota of *B. truncatus* across all sampling sites. Among the top 10 bacterial phyla identified in *B. truncatus*, two were also found in the aquatic bacterial communities: Cyanobacteria and Planctomycetota. Cyanobacteria was the second most abundant phylum in the water samples, while it ranked third among the top 10 phyla and seems less abundant in *B. truncatus*. Conversely Planctomycetota, which ranked seventh out of the 28 total phyla in the water samples, was significantly enriched in *B. truncatus* snails. This strengthens the idea that the snails may act as a narrower niche than aquatic environment selecting only some bacteria from the environment (Suzzi *et al*. 2023; Herlemann *et al*. 2024).

At the family level, the ‘common’ core microbiota of *B. truncatus* included Chitinophagaceae, Weekselaceae, and Flavobacteriaceae that all belong to the Bacteroidota phylum, as well as Comamonadaceae, Moraxellaceae, and Pseudomonadaceae belonging to the Proteobacteria phylum. Interestingly Proteobacteria, Bacteroidota and Firmicutes were previously detected as major bacterial taxa in a pool of *B. truncatus* initially originating from Spain and maintained in the laboratory for several generations (Huot *et al*. 2020). In particular, Flavobacteriaceae, Comamonadaceae and Pseudomonadaceae are families that are shared by this laboratory reared Spanish *B. truncatus* strain and all natural *B. truncatus* populations from northern Senegal studied here. These bacteria are likely to display important functions for the development of *B. truncatus*. In this regard, members of the Comamonadaceae family are known for their denitrifying activities and their action in the sulfur cycle (Khan *et al*. 2002; Wu *et al*. 2018). Pseudomonadaceae family, apart from containing pathogen bacteria for humans, can exhibit antimicrobial activity, help in nitrogen fixation and produce antifungal compounds (Fukuda *et al*. 2021; Saati-Santamaría *et al*. 2021). Pseudomonadaceae are also able to degrade all sort of complex molecular compounds including aromatic nucleus leading to break down environmental pollutants such as phthalates, hydroxybenzoates, and toluene (Parales and Harwood 1993; Przybylińska and Wyszkowski 2016).

Conversely, some bacterial families were found only in this study and were absent or present at low abundance in the previously studied *B. truncatus* laboratory strain. In particular, the Chitinophagaceae family, as their name implies, can hydrolyze chitin from the environment and are found primarily in soils and aquatic sediments (Lim *et al*. 2009; Madhaiyan *et al*. 2015). Under natural conditions, *B. truncatus* feed on recalcitrant food resources including detritus, decaying macrophytes, diatoms and filamentous algae, and thus potentially require a highly diverse microbial communities for digestion (Madsen 1992; Van Horn *et al*. 2012). In this regard, a reduction in the complexity of the food resource available for the laboratory population (i.e. organic washed lettuce) may also explain a relaxation of selection pressure and a loss of some bacteria taxa involved in snails’ digestion. Alternatively, although nonexclusively, the differences observed in the core microbiota between our *B. truncatus* field populations from Senegal and that of the Spanish laboratory strain may be a result of co-evolutionary processes driven in part by the evolutionary history of *B. truncatus* populations; the population established in Spain having probably diverged from the populations of Senegal for a long time. This latest hypothesis implies that the genetic structure of *B. truncatus* populations resulting from their evolutionary history influences the bacterial communities associated with these populations. This is precisely what our results show at the scale of northern Senegal, as we discuss in the next section.

### 2. Structuring factors of the microbiota of natural *B. truncatus* snails

Our result show that *B. truncatus* microbiota are well structured geographically although they display a less pronounced ‘distance–decay’ pattern than that of aquatic bacterial communities. Communities of free-living bacteria generally display such a pattern, although the robustness of the relationship may vary depending on the ecological context and the geographical scale (Clark *et al*. 2021). While the factors that lead to this ‘distance–decay’ pattern are still debated, it is accepted that it results at least from a limitation in the dispersion of bacterial communities and the heterogeneity of the environment (Green and Bohannan 2006; Sadeghi *et al*. 2021). We believe that, the strength of the ‘distance–decay’ pattern observed among aquatic bacterial communities in our study result from the fact that ecological heterogeneity and geographical distances between sites are tightly linked, with closer sites being associated to the same ecological system (e.g. lake de Guiers, River ‘Le Doué’). The less pronounced ‘distance-decay’ pattern of *B. truncatus* microbiota observed here is in line with the recent study from Herlemann *et al*. conducted on the freshwater snail *A. balthica* in Northern Europe at a similar geographical scale (Herlemann *et al*. 2024). As the authors suggest in their model, we can here hypothesize that *B. truncatus* create more uniform environments for their associated bacteria, compared to the heterogeneity of the aquatic environments where *B. truncatus* and aquatic bacterial communities are found. Another non-exclusive hypothesis would be that the structure of *B. truncatus* microbiota is determined by other predominant factors, and particularly some evolutionary factors specific to *B. truncatus* populations. In this respect, the MRM analysis shows that the genetic distance between *B. truncatus* hosts explains the beta diversity of their associated microbiota better (albeit slightly) than the geographical distance. This suggests that the evolutionary history of *B. truncatus* populations, which is strongly influenced by migration and drift as shown by the strong isolation by distance (IBD) pattern observed, is a key element in the structuring of *B. truncatus* microbiota. Thus, while phylosymbiosis between snail species and their microbiota is already documented at the inter-specific level (Huot *et al*. 2020; Schols *et al*. 2023), we here argue that phylosymbiosis could also occur at the intra-specific level.

The strong IBD pattern observed among *B. truncatus* populations was expected and fit well the results from previous genetic studies conducted on this species in different countries of Africa including Senegal. Early studies based on allozymes or a limited number of microsatellites have shown that self-fertilization is frequent in natural populations of *B. truncatus* (Njiokou *et al*. 1993; Jarne *et al*. 1994). On the other hand, populations of *B. truncatus* are often subjected to fluctuating episodes of extinction and recolonization that follow the alternation of wet and dry seasons that greatly shape the (sometimes temporary) habitats of this species. Episodes of extinction and recolonization by a small number of self-fertilizing individuals generally explain the low intra-site genetic diversity and genetic differentiation between populations, even if they are geographically close (Njiokou *et al*. 1993; Maes *et al*. 2022). Based on the 31 microsatellite markers newly developed in this study, we found no clear evidence of self-fertilization and the low values of genetic differentiation observed between geographically close populations suggest a non-negligible gene flow between these populations that tends to fade as the geographical distance between populations increases. We explain these results by the fact that all sites included in this study are permanent habitats with water present all along the year. This limits (but does not fully exclude) temporary population extinctions and allows populations to develop demographically, thus reducing the effect of genetic drift and potentially encouraging cross-fertilization. Moreover, water flow in the river Le Doué certainly promotes gene flow between geographically close populations and explains the little genetic differentiation observed between these populations. In this respect, it is interesting to note that the genetic markers used in this study reveal higher genetic differentiation between the population established in the Guia canal, which is fed and connected to the Le Doué River, compared to populations established along the Le Doué River at equal or greater geographical distances. Similarly, the sampling sites on the banks of Lake Guiers are only 3.2 km apart and it is likely that the sole effect of wind-generated surface currents facilitates gene flow between these two populations Conversely, high values of genetic differentiation were observed between geographically distant populations and are in line with the genetic patterns recently documented on *B. truncatus* populations in the same region based on 10750 SNPs (Maes *et al*. 2022).

Combined together, the genetic structure of *B. truncatus* populations and the geographical distance between populations do not fully explain the structure of the microbiota of *B. truncatus* and other factors, probably ecological, which we missed in this study, are certainly at work in shaping the microbiota of *B. truncatus*. For example, low genetic differentiation and little geographical distance clearly fail to explain the huge difference observed between the microbiota of *B. truncatus* established at the two sites along Lake Guiers (Saneinte and Mbane). Among the other potential structuring ecological factors that we investigated, the structure of aquatic bacterial communities does not predict the structure of *B. truncatus* microbiota when accounting for the genetic structure of *B. truncatus* populations and the geographical distances between sites. Thus, environmental bacterial communities do not directly influence *B. truncatus* microbiota. This was somewhat unexpected for at least two reasons. First, we might expect that some environmental bacteria are transferred to *B. truncatus* through several processes including ingestion during snail feeding or colonization of external snail tissues. Indeed, most freshwater snails are detritivorous or feed upon biofilms hence providing opportunity for different environmental bacteria to colonize and eventually establish within the gastrointestinal tract of snails established in different habitats (Madsen 1992; Van Horn *et al*. 2012; Kivistik *et al*. 2023). The lack of influence from aquatic bacterial communities on *B. truncatus* microbiota further supports the idea that *B. truncatus* acts as a filter for specific bacteria. This filtering effect, that might partly depend on the genetic background of *B. truncatus* lineages, limits the differences between the environmental bacterial communities that occasionally associate with *B. truncatus.* It is worth stressing that we here focused on *B. truncatus* whole microbiota and not specifically on the gut microbiota. By focusing only on these tissues, it is likely that we would have found a better congruence between environmental bacterial communities and *B. truncatus* gut microbiota. In fact, we did find some bacteria that are present both in the environment and in *B. truncatus* such as Chitinophagaceae as previously mentioned.

Finally, as part of the biotic environment, the presence of developing trematodes inside the snails does not seem to influence the snail microbiota although some infected *B. truncatus* displayed bacterial communities departing from those of non-infected sympatric congeners. In this respect, Portet *et al*. (2021) previously empirically showed that the experimental infection of laboratory-reared *Biomphalaria* snails by *Schistosoma mansoni* induced drastic modifications of snail microbiota soon after experimental infection, but that the microbiota gradually returns to its initial state a few days or weeks after. Consequently, it is likely that most *B. truncatus* diagnosed as infected with trematodes in our study had been infected for a sufficiently long time, allowing their microbiota to recover. Conversely, the rare infected *B. truncatus* displaying microbiota distinct from non-infected sympatric conspecifics could have been infected shortly before sampling. Alternatively, these individuals with different microbiota may have been exposed to a trematode but were incompatible from an immune perspective, potentially triggering an immune response that led to a temporary dysbiosis. Lastly, considering that we analyzed whole snail microbiota, we potentially sequenced the microbiota of the *B. truncatus* snail and its potential infecting parasites so we could find different microbial signatures depending on the infecting trematode species as observed by Salloum and collaborators in the mud snail *Zeacumantus subacarinatus* infected by four different trematode species (Salloum et al. 2023). However, due to the low observed prevalence of infection in our study, we might have insufficient results to robustly explore the link between the infection status and the microbiota composition of *B. truncatus* which means that we may have missed such signals.

### 3. Conclusions

Our study highlights a well-structured geographical pattern in the microbiota of natural populations of *B. truncatu*s that is best explained by the genetic structure of *B. truncatus* populations. This suggests an important role of the evolutionary trajectory of *B. truncatus* populations in shaping their microbiota. This calls for further studies to identify the potentially genomic determinants that influence the composition of hosts microbiota. Importantly however, some strong differences in the composition of *B. truncatus* microbiota from genetically and geographically close populations remain unexplained and suggest that some unexplored ecological factors or hosts’ physiological traits (e.g. age or fitness) could also influence the microbiota of this species. Among these ecological factors we found that neither the bacterial communities from the surrounding aquatic environment nor the presence of trematode developing within hosts influence the overall structure of hosts microbiota. The temporal monitoring of the microbiota associated with hosts populations and that of the environment in which the hosts are established could provide a valuable approach to identify potential environmental factors at play in shaping host microbiota. In the case of *B. truncatus*, populations from habitats with high temporal environmental fluctuations such as temporary ponds should be targeted. More generally, such temporal monitoring studies in more temperate regions with important seasonal environmental changes would help better understanding the potential environmental factors influencing the microbiota of freshwater gastropods. Finally, further research on the interactions between genetic and environmental variables would be necessary to specifically assess the plasticity of hosts microbiota hence contributing to a better understanding of the relationship between the environment, hosts and their microbiota at the intra-specific level.

## 4. Acknowledgements

We warmly thank the Bio-Environment platform (UPVD, Région Occitanie, CPER 2007-2013 Technoviv, CPER 2015-2020 Technoviv2) and Jean-François Allienne, Margot Doberva and Michèle Laudié for support in library preparation and sequencing. Microsatellites development and genotyping were performed at the PGTB (doi:10.15454/1.5572396583599417E12) with the help of Zoé Compagnie, Adline Delcamp and Erwan Guichoux. This study was supported in part by the European and Developing Countries Clinical Trials Partnership (EDCTP2) program (TMA2018CDF-2370), supported by the European Union. It was funded by the French Agency for Food, Environmental and Occupational Health & Safety (PNRES 2019/1/059 Molrisk) and the Occitanie Region (Schistodiag program). This study was carried out with the support of LabEx CeMEB, an ANR ‘Investissements d’avenir’ program (ANR-10-LABX-04-01), and within the framework of the ‘Laboratoire d’Excellence (LABEX)’ TULIP (ANR-10LABX-41). It was also supported by the MICROVECT Project (défi clé RIVOC Occitanie Region, University of Montpellier).

## Data availability statement

All sequence data generated have been submitted to the Sequence Read Archive of NCBI.

## Ethics approval statement

The project has received approval from the National Ethical Committee (CNERS) of Senegal (agreement number: 00061/MSAS/CNERS/SP).

## 6. Data Accessibility and Benefit-Sharing

### 6.1. Data Accessibility Statement

All sequence data generated have been submitted to the Sequence Read Archive of NCBI.

### 6.2. Benefit-Sharing Statement

This project obtained from the Nagoya office in Senegal, an exemption from authorization of access and use of genetic resources (number: 001339 of November 15, 2021, reference: V/L du 28 octobre 2021) by the Competent National Authority (Directorate of National Parks of Senegal).

## 7. Authors Contributions

O.R., P.D., B.G., M.J. were responsible for the research design. O.R., P.D., B.S., J.B. conducted the field work. M.J. and P.D. conducted the lab work. O.L. and E.C. developed the microsatellite dataset. M.J., P.D., O.R., E.T., and T.L. contributed to the data analysis. M.J., O.R. and P.D. wrote the initial draft of the manuscript and all authors contributed to revisions. O.R. and B.G. supervised the project.

