## Supplementary material for "Evidences that host genetic background more than the environment shapes the microbiota of the snail *Bulinus truncatus*, an intermediate host of *Schistosoma* species": Suppl. Fig.

Table of Contents:


| Appendix number | Title | Page |
|---|---|---|
| 1 | Infection status of B. truncatus snails | 2 |
| 2 | Microsatellite development | 2 |
| 3 | Sequence-Based Microsatellite Genotyping | 3 |

| Fig number | Title | Page |
|---|---|---|
| Suppl. fig 1 | Scatterplot displaying Nei's genetic distance between individuals as a function of the geographic distance across sampling sites. | 7 |
| Suppl. fig 2 | Alpha diversity computed on the bacterial communities from the environmental water-sediment samples and isolated from *B. truncatus* snail hosts at each sampling site. | 7 |
| Suppl. fig 3 | Alpha diversity computed on the bacterial communities isolated from *B. truncatus* snail hosts at each sampling site. | 8 |
| Suppl. fig 4 | Relative abundance of Cyanobacteria and Plantomycetota phyla in environmental samples and *B. truncatus* snails among the different sampling sites. | 9 |
| Suppl. fig 5 | Principal coordinate analysis (PCoA) plot showing environmental samples and *B. truncatus* snail associated bacterial communities distributed along the two first PCoA axes computed based on pairwise Bray-Curtis distance obtained between each sample. | 9 |
| Suppl. fig 6 | Principal coordinate analysis (PCoA) plot showing the bacterial communities of *B. truncatus* snails according to their original sampling sites, and to their infectious status distributed along the two first PCoA axes based on pairwise Bray-Curtis distance computed between each sample. | 10 |

| Table number | Title | Page |
|---|---|---|
| Suppl. table 1 | Abundance of *B. truncatus*, trematode prevalence among sites. | 4 |
| Suppl. table 5 | Results of Kruskall-Wall test comparing the *B. truncatus* alpha-diversity depending on the sampling site. | 8 |
| Suppl. table 6 | Results of Kruskall-Wall test comparing the *B. truncatus* alpha-diversity depending on the infectious status of individuals. | 8 |
| Suppl. table 7 | Results of PERMANOVA analysis comparing the *B. truncatus* beta-diversity based on Bray-Curtis distance revealing significant differences depending on the sampling site but not on the infectious status of the hosts. | 10 |

**Appendix 1:** Infection status of *B. truncatus* snails

The presence of developing trematodes within each of the 124 snails was diagnosed using the Trem_16S_F1 (GACGGAAAGACCCCRAGA) and Trem-16S_R2 (CRCCGGTYTTAACTCARYTCAT) *16S* trematode metabarcode developed by (Douchet *et al.* 2022). DNA extract from each snail was first diluted to 1/100$^{th}$ to limit the potential effect of PCR inhibitors and hence limit PCR false negatives. PCRs were performed using the GoTaq® G2 Hot Start Polymerase kit (Promega). Each PCR reaction contained Green Buffer at 1X, $MgCl_2$ at 1.5mM, dNTPs at 0.2µM, forward and reverse primers at 0.4µM, 1.25 units of GoTaq G2 Hot Start, 2µL of 1/100 DNA (~ 1.5 ng) sample and ultrapure water in a final reaction volume of 10µL. The PCR program was as follows: an initial denaturation step at 94°C for 3 min, followed by 40 cycles of denaturation at 95°C for 30 sec, hybridization at 54°C for 30 sec and elongation at 72°C for 15 sec, and a final elongation step at 72°C for 5 min. A subset of 5µL of the resulting PCR products were migrated on a 2% agarose gel for 30 min at 110V and revealed under UV. Positive samples were reamplified in a final volume of 25µL using the same Trem_16S_F1 and Trem_16S_R2 primers implemented with Illumina sequencing adapters and clear Buffer.

**Appendix 2:** Microsatellite development

The *B. truncatus* genome (GenBank accession GCA_021962125.1, (Young *et al.* 2022)) was used for microsatellite discovery using the software QDD v3.1.2 (Meglécz *et al.* 2010). Primer design parameters were set to target 100 pb to 180 pb amplicons and with primer parameters optimized for multiplex PCR (Lepais *et al.* 2020). A total of 96 primer pairs (9 tetranucleotide and 66 trinucleotide motifs with more than 6 repeats, and 21 dinucleotide motif with more than 16 repeats) were selected based on criteria maximizing amplification success (Meglécz *et al.* 2014) and polymorphisms (i.e. high number of repeats). PrimerPooler software (Brown *et al.* 2017) was used to determine two sets of markers to co-amplify into two multiplex PCR (W1 and W2**, suppl. Table 2**). The primers were built by adding the universal sequence ACACTCTTTCCCTACACGACGCTCTTCCGATCT to the 5' end of each forward primer and the universal sequence GTGACTGGAGTTCAGACGTGTGCTCTTCCGATCT at the 5' end of each reverse primer.

Each pair of primers was tested for amplification on a DNA pool from a subset of *B. truncatus* specimens. The PCR was performed in a final volume of 10 µL using Hot Firepol Blend master mix (Solis Biodyne), 10 ng of DNA and 0.2 µM of each primer. The PCR conditions consisted of an initial denaturation at 95°C for 15 min followed by 35 cycles of denaturation at 95°C for 20 sec, annealing at 59°C for 60 sec, extension at 72°C for 30 sec, and a final extension step at 72°C for 10 min. Amplification was checked on a 3% agarose gel. Primer pairs that showed consistent amplification with amplicon at the expected size were kept for multiplex PCRs.

**Appendix 3:** Sequence-Based Microsatellite Genotyping

Each multiplex PCR amplification was performed in a final volume of 5 µL using 5X Hot Firepol Multiplex master mix (Solis Biodyne), 0.05 µM of each primer, and 10 ng of DNA. The PCR conditions consisted of an initial denaturation at 95°C for 12 min followed by 35 cycles of denaturation at 95°C for 30 sec, annealing at 59°C for 180 sec, extension at 72°C for 30 sec, and a final extension step at 72°C for 10 min. The sequencing libraries of each multiplex were constructed using a second PCR that attached adapters and sample-specific pairs of indices (10 bp unique sequences) to each side of the amplicons by targeting the universal sequence attached to the locus specific primers. The indexing PCR is set up in a volume of 5 µL using 5X Hot Firepol Multiplex master mix (Solis Biodyne), 1.25 µL of amplicon and 0.5 µM of each of the forward and reverse adapters. The PCR conditions consisted in an initial denaturation at 95°C for 12 min followed by 15 cycles of denaturation at 95°C for 30 s, annealing at 59°C for 90 s, extension at 72°C for 30 s, and a final extension step at 72°C for 10 min. Libraries for each multiplex were then pooled, purified with 1.2× *SPRI magnetic beads* and sized using the Pippin Prep System (Sage Science; 2% agarose, 200-450 bp). We checked the library quality on a Tapestation 4200 (Agilent Technologies, Santa Clara, CA) and quantified it using QIAseq Library Quant Assay kit (Qiagen) on a Mic qPCR Cycler (Bio Molecular Systems). An equimolar pool of the two multiplexes (W1 and W2) was finally sequenced on an Illumina iSeq100 sequencer with a 2 × 150 bp paired-end sequencing kit. A random subsample of 95 samples was genotyped twice to optimize the bioinformatic pipeline to each locus, estimate locus-level allelic error rate (number of allele mismatches between replicates divided by the total number of alleles compared), and finally select loci that produced repeatable genotypes for the final genotypic dataset.

The bioinformatics analysis consisted in three steps: (1) sequence preparation and merging of sequence pairs with BBmerge v38.87 (Bushnell et al. 2017), (2) preliminary marker validation by comparing replicates and testing different bioinformatic parameters and (3) final genotyping using the optimal parameters on the validated markers.

Genotype calling from allele sequence was performed using FDSTools (v1.2.0; (Hoogenboom et al. 2017)) embedded in a pipeline that compares the replicate genotypes to compute allelic error rate and format the results and associated locus and allele information (see (Lepais et al. 2020) for more details, documented pipeline available here: https://data.inra.fr/dataset.xhtml?persistentId=doi:10.15454/HBXKVA). Sequence coverage of each allele was used to estimate allele dosage and determine complete tetraploid genotypes following (Cui et al. 2022). Microsatellite markers monomorphic across all individuals were removed from the dataset as well as those that display an excess of missing data over individual genotypes. This led to the selection of a final set of 31 microsatellite markers. We also removed individuals that display missing data, at least, one of the 31 microsatellites. After this filtering process, our genetic database consisted in a total of 110 individuals (over the 124 initially genotyped snails) fully genotyped at 31 microsatellite markers.

**Suppl. table 1**: Abundance of *B. truncatus*, trematode prevalence among sites

| Locality | Habitat | X coordinate | Y coordinate | Sampling date | Number of shell extracted Bulinus | Trematode prevalence among the diagnosed shell extracted snails | Number of trematodes species | Volume of filtered water (L) |
|---|---|---|---|---|---|---|---|---|
| Ndiawara | River 1 | 16°35′04″N | 14°50′58″W | 11/02/2022 | 8 | 25% | Paramphistomoidea 1 (n=1)<br>Paramphistomoidea 2 (n=1)<br>*Petasiger sp.* (n=1) | 6,5 |
| Ouali Diala | River 1 | 16°35′56″N | 14°56′10″W | 11/02/2022 | 12 | 25% | Paramphistomoidea 1 (n=3)<br>Paramphistomoidea 2 (n=3)<br>*S. haematobium* (n=1) | 10 |
| Dioundou | River 1 | 16°35′50″N | 14°53′22″W | 12/02/2022 | 5 | 0 | NA | 10 |
| Fonde Ass | River 1 | 16°36′20″N | 14°57′38″W | 12/02/2022 | 17 | 29% | *Haematolochus sp.* (n=5) | 10 |
| Khodit | River 1 | 16°35′45″N | 14°56′42″W | 13/02/2022 | 16 | 25% | *S. haematobium* (n=4)<br>*S. bovis* (n=1)<br>*Haematolochus sp.* (n=1) | 5 |

| | | | | | | | | | |
|---|---|---|---|---|---|---|---|---|---|
| Guia | Inlet | 16°35′51″N | 14°55′31″W | 12/02/2022 | 14/02/2022 | 17 | 5.8% | S. *haematobium* (n=1)<br>*Haematolochus sp.* (n=1) | 5 |
| Mbane | Lake | 16°16'15"N | 15°48'7"W | 15/02/2022 | | 16 | 0 | NA | 2.8 |
| Saneinte | Lake | 16°14'32"N | 15°48'6"W | 15/02/2022 | | 17 | 0 | NA | 3.5 |
| Lampsar | River 2 | 16°6'34"N | 16°20'58"W | 17/02/2022 | | 16 | 25% | *Orientocreadium batrachoides* (n=4)<br>*S. bovis* (n=1) | 18 |

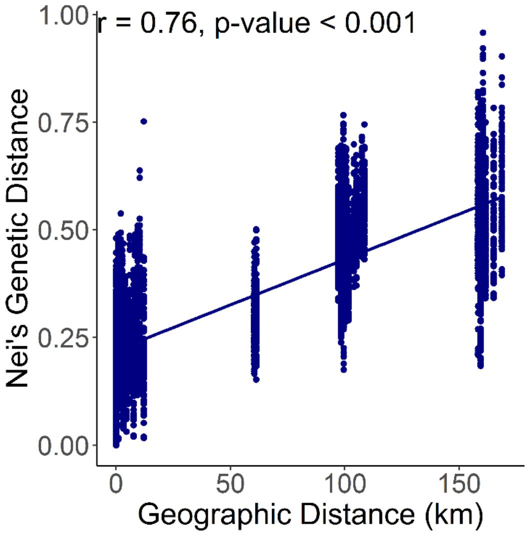


**Suppl. fig 1:** Scatterplot displaying Nei's genetic distance between individuals on 31 loci as a function of the geographic distance across sampling sites and showing a strong isolation by distance pattern.

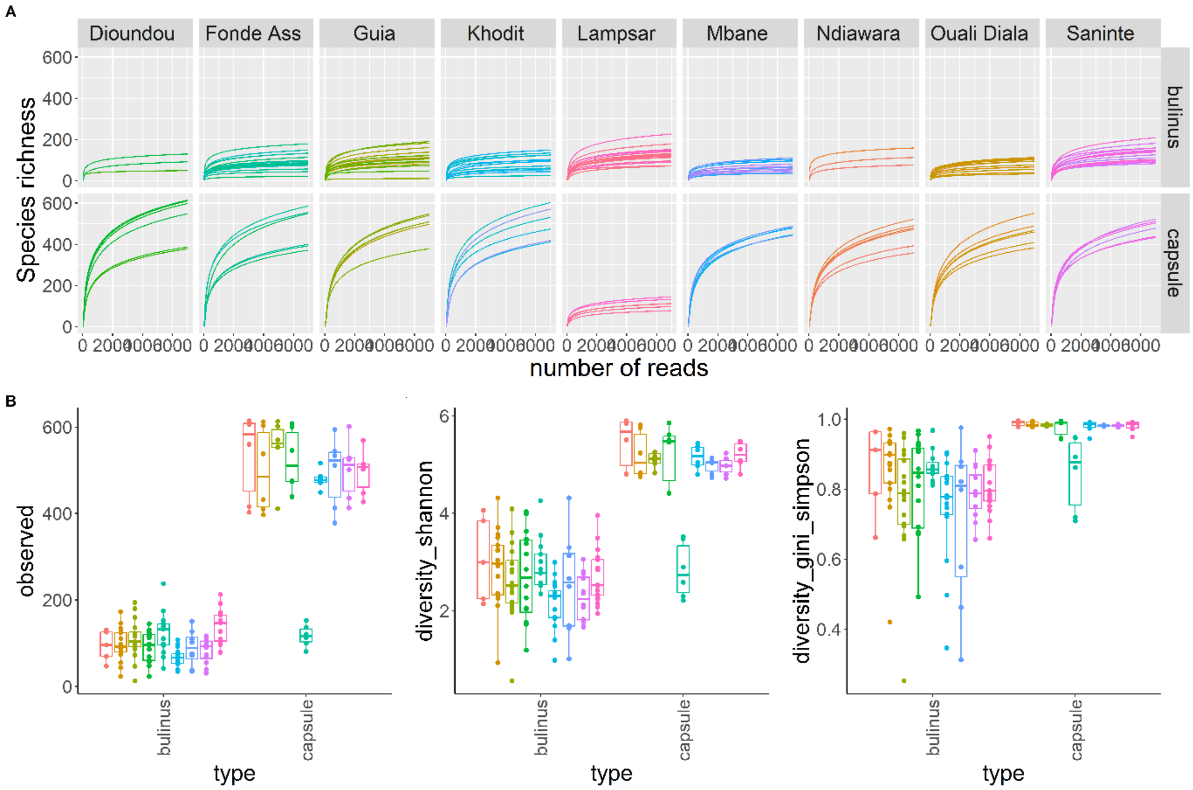


**Suppl. fig 2**: Alpha diversity computed on the bacterial communities from the environmental water-sediment samples and isolated from *B. truncatus* snail hosts at each sampling site. **(A)** Rarefaction curves computed per sites and sample types (e.g. *B. truncatus* snails versus environmental samples) showing that all samples reached saturation at the chosen rarefaction threshold. **(B)** Indices of microbial alpha diversity as a function of sample type (e.g. *B. truncatus* snails versus environmental samples) and sampling sites were significant when using observed ASV richness, Shannon's diversity index and Simpson's diversity index.

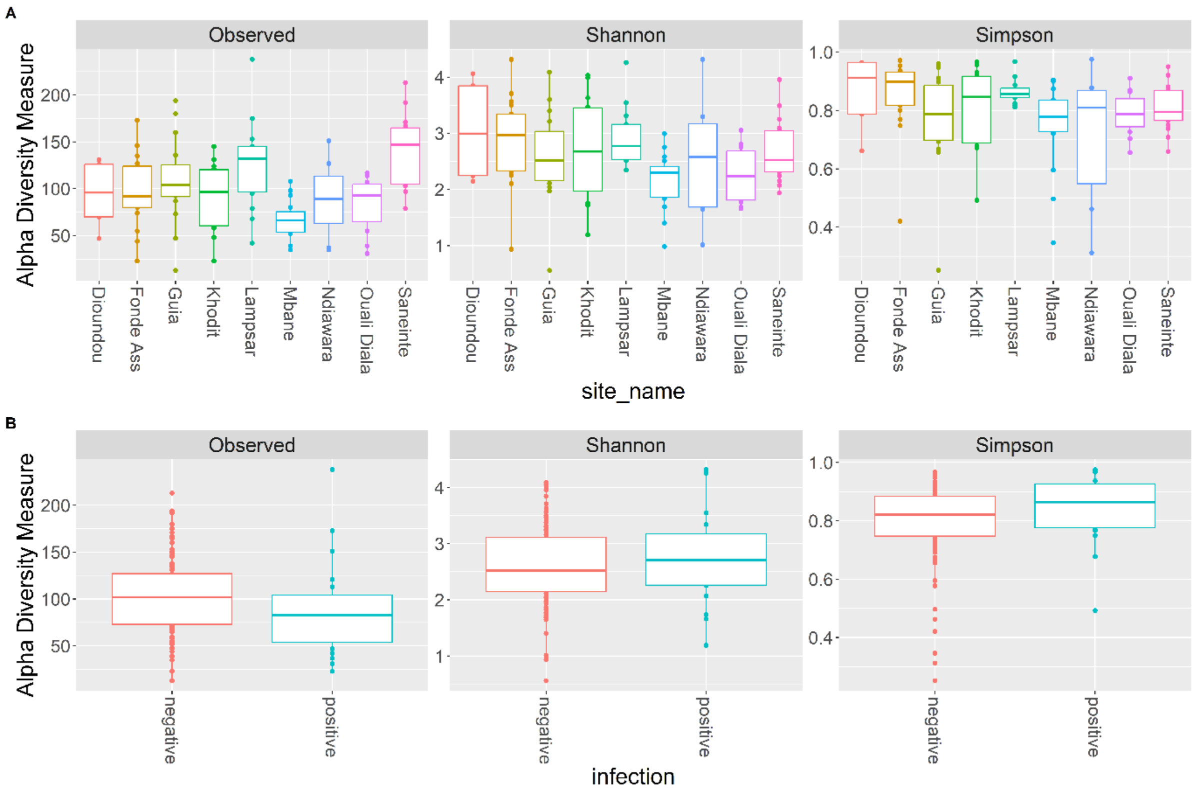


**Suppl. fig 3**: Alpha diversity computed on the bacterial communities isolated from *B. truncatus* snail hosts at each sampling site. Comparisons of alpha diversity indexes computed on snail host microbiota between snails hosts among sampling sites (**A**) and depending on the infectious status (**B**) using observed ASV richness, Shannon's diversity index and Simpson's diversity index.

**Suppl. table 5:** Results of Kruskall-Wall test comparing the *B. truncatus* alpha-diversity depending on the sampling site of individuals based on ASV richness, diversity Shannon and Gini-Simpson indices revealing significant differences (all p-value < 0.05). Df = degree of freedom.

| | Kruskal-Wallis Chi Squared | Df | p-value |
|---|---|---|---|
| ASV richness | 32.536 | 8 | **< 0.05** |
| Diversity Shannon | 17.82 | 8 | **0.02262** |
| Diversity Gini-Simpson | 16.109 | 8 | **0.04085** |

**Suppl. table 6:** Results of Kruskall-Wall test comparing the *B. truncatus* alpha-diversity depending on the infectious status of individuals based on ASV richness, diversity Shannon, and Gini-Simpson indices revealing no significant differences (all p-value > 0.05). Df = degree of freedom.

| | Kruskal-Wallis Chi Squared | Df | p-value |
|---|---|---|---|
| ASV richness | 2.7537 | 1 | 0.09703 |
| Diversity Shannon | 0.39046 | 1 | 0.5321 |
| Diversity Gini-Simpson | 2.1672 | 1 | 0.141 |
| Pielou index | 2.209 | 1 | 0.1372 |

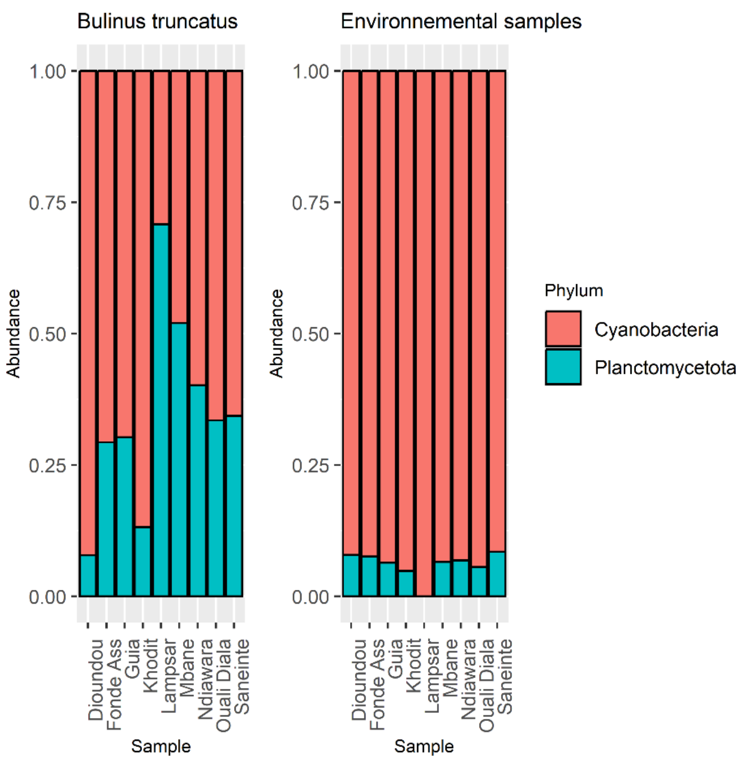


**Suppl. Fig 4:** Relative abundance of Cyanobacteria and Plantomycetota phyla in environmental samples and *B. truncatus* snails among the different sampling sites.

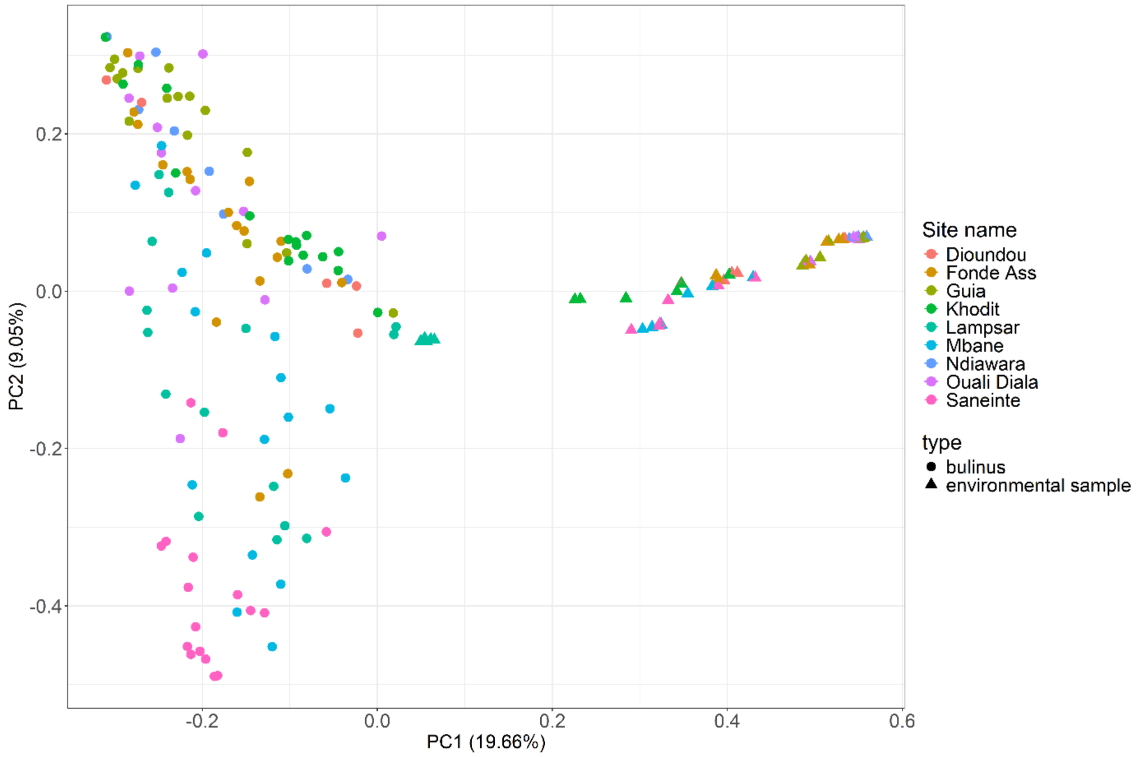


**Suppl. fig 5**: Principal coordinate analysis (PCoA) plot showing environmental samples (triangles) and *B. truncatus* snail associated (dots) bacterial communities distributed along the two first PCoA axes computed based on pairwise Bray-Curtis distance obtained between each sample.

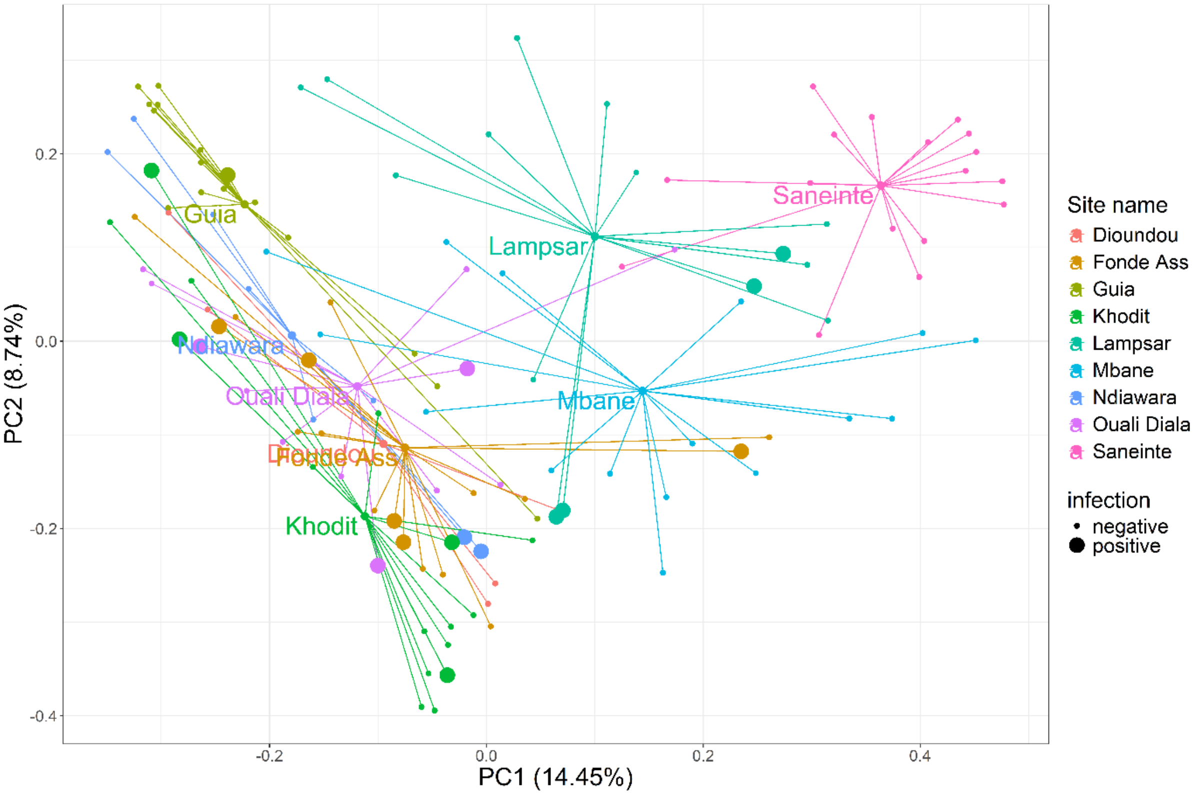


**Suppl. fig 6**: Principal component analysis (PCoA) plot showing the bacterial communities of *B. truncatus* snails according to their original sampling sites (see color legend), and to their infectious status (dot size) distributed along the two first PCoA axes based on pairwise Bray-Curtis distance computed between each sample. Each colored small dot represents one snail host from a given sampling site. Large dots are snail hosts positive to trematode molecular diagnosis. Medium dots are the centroids of the bacterial communities of snails originating from each sampling site.

**Suppl. table 7:** Results of PERMANOVA analysis comparing the *B. truncatus* beta-diversity based on Bray-Curtis distance revealing significant differences depending on the sampling site but not on the infectious status of the hosts. Df = degree of freedom, F = F-statistic. The significant values are in bold.

| | Df | Sum of squares | $R^2$ | F | P-value |
|---|---|---|---|---|---|
| Site | 1 | 3.583 | 0.09184 | 12.197 | **< 0.05** |
| Infection | 1 | 0.384 | 0.00984 | 1.307 | 0.1263 |
| Site:infection | 1 | 0.380 | 0.00973 | 1.292 | 0.1359 |
